# Impact of charge patches on tumor disposition and biodistribution of therapeutic antibodies

**DOI:** 10.1101/2021.09.01.458024

**Authors:** Jakob C. Stüber, Karin F. Rechberger, Saša M. Miladinović, Thomas Pöschinger, Tamara Zimmermann, Remi Villenave, Miro J. Eigenmann, Thomas E. Kraft, Dhaval K. Shah, Hubert Kettenberger, Wolfgang F. Richter

## Abstract

This study explores the impact of antibody surface charge on tissue distribution into various tissues including tumor. Tumor-bearing mice were dosed intravenously with a mixture comprising three antibodies engineered to carry negative charge patches, a balanced charge distribution, or positive patches, respectively. Tissue levels were analyzed with a specific LC-MS/MS method. In addition, the antibody mix was administered to non-tumor bearing mice. Muscle and skin interstitial fluid were obtained by centrifugation and analyzed by LC-MS/MS. An in-vitro endothelium model was explored for its feasibility to mimic the observed distribution differences. A balanced charge distribution was optimal in terms of total tumor exposure, while in other tissues negatively charged and balanced charged antibodies gave similar results. In contrast, positive charge patches generally result in increased serum clearance but markedly enhance tumor and organ uptake, leading to higher tissue-to-serum ratios. The uptake and availability in the interstitial space were confirmed by specific assessment of antibody levels in the interstitial fluid of muscle and skin, with similar charge impact as in total tissue. The in vitro model was able to differentiate the transport propensity of this series of antibody variants. In summary, our results show the differential effects of charge patches on an antibody surface on biodistribution and tumor uptake. These insights may help in the design of molecules with biodistribution properties tailored to their purpose and an optimized safety profile.

## Introduction

Monoclonal antibodies have gained tremendous importance in the treatment of a wide range of diseases over the last two decades [1]. The distinctive combination of high target binding affinity, exquisite specificity, and extraordinarily long plasma half-lives have been pivotal to their clinical success [2]. The exceptional plasma residence time is afforded by salvage mediated through the neonatal Fc receptor (FcRn), which protects antibodies from rapid lysosomal degradation [2,3]. The FcRn interaction also affects biodistribution and tissue catabolism of antibodies [4–6].

Arguably, it is the delivery to the target site, which restricts efficacy of antibody therapy in many cases. In particular, improving uptake into solid tumors is an important challenge for current research [7]. The presence of cells exposed to insufficient amounts of antibody within a tumor under treatment not only restricts efficacy, but may also foster the development of resistance against a targeted treatment [8].

Following intravenous injection, the most frequent route of administration for therapeutic antibodies [9], sustained high plasma concentrations drive peripheral tissue and tumor uptake (Figure 1a, [3,8,10]). The site of action for most antibody therapeutics of solid tumors is the cellular surface, accessed from the interstitial space [7]. Several barriers need to be crossed to reach this compartment and distribute homogeneously (Figure 1a, [8,11]).

**1.**
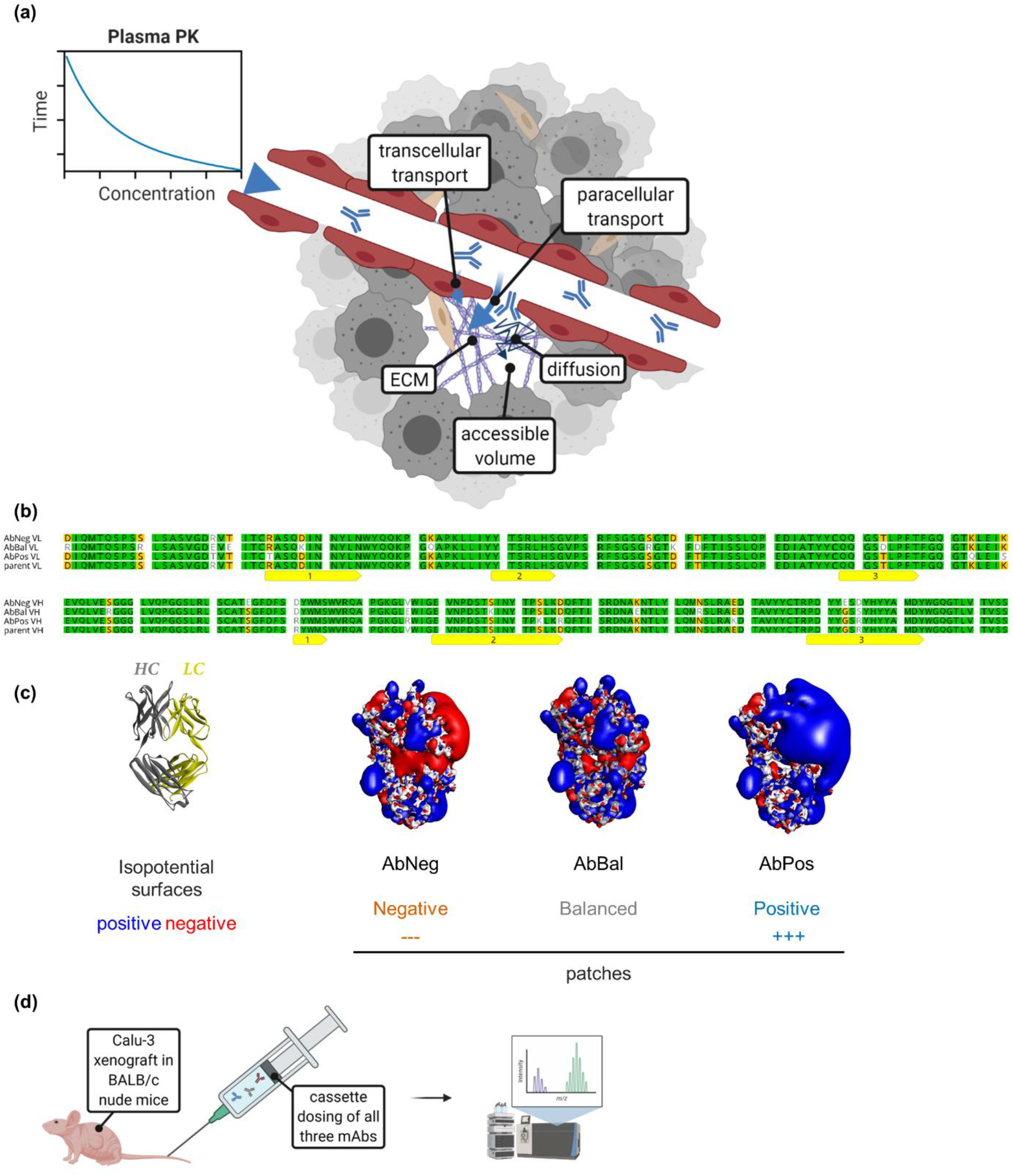
Model of tumor uptake, used antibody variants and study outline: (a) Processes governing tumor uptake of antibodies and potential impact of surface charge patches. (b) Isocontour representation of the three charge-patch variants used in the study. (c) Sequence alignment of the designed variable regions, with the complementarity-determining regions (CDRs) highlighted in yellow. (d) Design and readout of the in vivo xenograft study.

As a first step of tissue uptake, antibodies need to traverse the capillary wall. It is commonly assumed that the major extravasation route in tumors is paracellular, because tumor capillaries were found to be highly irregular and leaky due to fenestrae and transcellular holes [12,13]. At the same time, however, poor lymphatic drainage leads to an elevated interstitial pressure within tumors, which impedes convective influx and results in a lower net antibody uptake. Thus, paracellular tumor uptake mostly relies on diffusion [8]. Alternatively, transcellular transport (i.e., transcytosis) may contribute to the net tumor uptake, in particular if it is enhanced through binding to FcRn or another internalizing receptor expressed in endothelial cells. It is generally assumed that extravasation is the limiting step for tumor uptake [2,8].

After passing the capillary wall, antibodies need to migrate through the tumor interstitial space by diffusive transport. A high local concentration of free binding sites can effectively limit antibody tumor penetration to a narrow space around the blood vessels, a phenomenon termed “binding site barrier” [14]. This is exacerbated by high antigen expression and turnover [8], and in some cases, also the mode of action, which may involve modulation of the target turnover. In addition, biophysical properties can influence diffusive transport in the interstitial space. For example, nonspecific, usually charge-based interactions with high-abundance binding sites formed by the extracellular matrix or the cell surface (glycocalyx and membrane constituents) may retard diffusion [15]. Finally, the accessible volume, the fraction of the tumor interstitial space that can actually be occupied by antibody molecules, which depends strongly on molecular size [16], is a major determinant of the total antibody uptake.

Charge-related biophysical properties, such as isoelectric point (pI), overall charge, and charge distribution (patches), are known to strongly influence antibody pharmacokinetics and biodistribution [17]. Numerous studies have shown that cationization generally results in significantly increased plasma clearance, likely because of increased charge-mediated endothelial uptake [17–19]. This is accompanied by higher (apparent) volumes of distribution, and increased peak organ concentrations. Conversely, anionization or neutralization (of positive charges) consistently reduces tissue uptake and often, at least if the modifications are introduced through genetic engineering and not through chemical conjugation, lowers clearance [17–21].

Charge-related effects can conceivably affect several of the processes involved in antibody tumor disposition (Figure 1a). Some studies found a strong impact of the extracellular matrix (ECM) on interstitial antibody diffusion [22–26]. This suggests the presence of charge-mediated interactions with ECM components, e.g. polyanionic glycosaminoglycans, which may differ depending on antibody charge properties. Also, diffusion-limited uptake routes, i.e. paracellular transcapillary transport in the leaky tumor vasculature [8,27] and surface transport [11], could be hampered by charge-dependent transient binding events. Transcellular extravasation has also been shown to be modulated by charge, through promotion of charge-mediated adsorptive transcytosis [28]. Finally, overall isoelectric point (pI) was found to influence total tumor uptake at steady state due to differences in the accessible volume fraction, i.e. the part of the tumor space that can actually be occupied by a molecule [29]. However, a rather robust proportionality between plasma and tissue concentration was observed for a dataset containing numerous biodistribution studies from several species. This suggested that tissue-to-serum ratios would not differ strongly among molecules, although biases in selection of the molecules (for properties desirable in drug development) cannot be excluded [30]. Thus, charge properties are generally believed to modulate vascular permeability [17,27] and slightly alter the steady-state levels by impacting the accessible volume, but not to effect a significant increase in uptake or tissue-to-plasma exposure ratios.

This study focused on the impact of the surface charge distribution on tumor disposition and tissue distribution in other tissues. Surface charge distribution, i.e. the presence of charge patches, is a biophysical property, which can be engineered in an antigen-independent manner. We hypothesized that small modifications of a single common ancestor molecule would provide a defined system to study charge-related effects. Using protein engineering, we covered a wide range of possible charge distribution profiles. These variants were then studied with respect to their tumor uptake and biodistribution in other tissues, with a detailed analysis of specific exposure profiles. In addition, the actual interstitial levels of the substances delivered to tissues were experimentally determined. Finally, we developed an *in vitro* assay, which is predictive of *in vivo* extravasation properties, and enables mechanistic insight into the charge-patch dependent mechanisms of tissue uptake.

## Materials & Methods

### Design of the surface-charge distribution variants

Starting from an existing human IgG1 antibody, which was engineered to abolish binding to its target, human and murine CD44, we manually identified surface-exposed residues in the variable domain for charge-patch engineering. Specifically, residues were selected whose side chains do not interact with other parts of the variable domain and which were in spatial proximity (<10 Ångström) to one another. Mutation sites were both CDR and framework residues. Sequence details are shown in Figure 1c. Isocontour renderings shown in Figure 1 were generated using PyMOL (Schrödinger, LLC)

### Protein preparation and analytics

Antibodies were prepared in-house according to standard procedures, using transient transfection in HEK293 cells, capture on a Protein A column, and preparative size exclusion chromatography. All samples used in animal experiments were verified to show a monomer content >95% in size exclusion chromatography, >95% purity in non-reducing capillary electrophoresis in the presence of SDS, <0.200 EU/ml endotoxin content (protein concentration ≥5 mg/ml), and the identity of their protein content was confirmed by electrospray ionization – mass spectroscopy (ESI-MS).

### Xenograft in-life experiment for tumor disposition and biodistribution

Female Balb/c nude mice (Charles River, Sulzfeld, Germany) were subcutaneously inoculated with 5×10^6^ cells of the human lung adenocarcinoma cell line Calu-3 (HTB-55, ATCC, VA, USA) in 100 µL phosphate-buffered saline (PBS). At a tumor volume of 100-150 mm^3^, animals were randomized into groups of n=3 animals per group. All animals received a single intravenous dose of an antibody mixture containing the three variants of non-binding anti-CD44 antibodies (5 mg/kg each) with differing charge-related biophysical properties (AbNeg, AbBal, AbPos). Animals were sacrificed at 3, 6, 24, and 48 h post-dose (n=3/time point) and tissues of interest (tumor, muscle, liver, and spleen) were collected and weighed before storage at −20°C until analysis. Five minutes prior to sacrifice, animals received an IV bolus injection of infliximab at 5 mg/kg to be used as a vascular marker. The cell line was confirmed to be free of murine pathogens and murine viruses (Biomedical Diagnostics, Hannover, Germany). The animal experiment was performed in accordance with the guidelines stated by the Federation for Laboratory Animal Science Associations (FELASA) and the applicable national animal welfare law. Animal experiments were approved by the local Government of Upper Bavaria ethics committee (Regierung von Oberbayern, Munich, Germany) and performed under license ROB-55.2.- 2532.Vet_02-19-5. All animals were kept under specific pathogen free (SPF) conditions in an animal facility accredited by the Association for Assessment and Accreditation of Laboratory Animal Care (AAALAC).

### In-life experiment for serum pharmacokinetics and interstitial fluid isolation from muscle and skin

C57BL/6 mice (Charles River Laboratories, Lyon, France) received a single intravenous dose of a mixture containing three variants of anti-CD44-antibodies (5 mg/kg each). Animals were sacrificed at 3, 5.5, 24, 48, and 96 h post-dose (n=3/time point) and blood, muscle and skin were collected. 5 min prior to sacrifice, animals received an IV bolus injection of infliximab at 5 mg/kg to be used as a vascular marker. Blood was allowed to clot and serum prepared. Muscle and skin interstitial fluid (ISF) was prepared by low speed centrifugation as described elsewhere [31]. In brief, freshly harvested tissue was placed on a Nylon mesh inserted in an Eppendorf tube and centrifuged for 10 min at 450 x g. Collected interstitial fluid and serum was stored frozen until analysis. The animal experiment was carried out with permission of the Veterinary Office of Canton Basel City, in accordance with the Swiss regulations, in an animal facility accredited by the AAALAC.

### Bioanalytics of serum, tissues and interstitial fluid

The bioanalysis of antibodies was conducted using bottom-up LC-MS/MS approach with immunoaffinity-enrichment utilizing protein A conjugated magnetic beads.

### Sample pre-treatment

Tissue samples were homogenized in a 4-fold volume of 100 mM PBS solution (pH=7.4). Serum (30 µL), interstitial fluid (15 µL) or tissue homogenate (200 µL) were separately added to PBS-washed protein A conjugated magnetic beads. A generic, stable-isotope-labeled monoclonal antibody (SILu™Mab, Sigma) was then added as an internal standard, and the plate was incubated for 90 min.

After the supernatant was discarded, captured antibodies were eluted with 0.1% trifluoroacetic acid (TFA, 150 µL) and neutralized with 1 M Tris-HCl (50 µL). Enriched sample fractions were then digested at 70 °C for 90 min using a SMART Digest™ Trypsin Kit (Thermo Scientific, 5-10 ng of trypsin/sample) and acidified with formic acid (2 µL). In order to facilitate sample processing and increase analytical throughput, all steps (including immuno-affinity enrichment and tryptic digestion) were performed in the 96-well format.

### LC-MS/MS

For LC-MS analysis, 30 µL of each digested sample was injected onto the analytical column (Aeris peptide XB-C18, 2.1×150mm, 2.6 µm), the signature proteotypic peptides were separated from matrix interferences using a gradient elution from 0.1% formic acid to 30% acetonitrile and detection performed in multiple reaction monitoring (MRM) mode using a triple quadrupole mass spectrometer (Sciex QTrap 6500).

The calibration range covered 3 orders of magnitude. The precision and accuracy of the method successfully met all pre-defined acceptance criteria for this study. A minor interfering signal in muscle samples was observed for AbNeg, and calibrators were thus corrected for pretest levels.

### Data processing and analysis

#### Exposure analysis

Noncompartmental analysis of serum data was performed using MATLAB R2020a and SimBiology (The MathWorks, Inc.), applying the option for sparse data. Similarly, the area under the time-concentration curve and the associated variance for the tissues was calculated according to Nedelman & Jia [32] with the modification of Holder [33].

#### Antibody biodistribution coefficients

Antibody biodistribution coefficients (ABC) were calculated following [30], however only based on the 48-h-time point, to exclude any effect of differing equilibration kinetics. Tissue levels were obtained from our biodistribution study in BALB/c nude mice, and serum levels from CL57/BL6 mice (with equal dosing) to calculate ABCs.

#### Residual plasma correction

Residual plasma correction was based on the co-injection of a non-interfering antibody (infliximab) as a vascular marker 5 min prior to sacrifice (to minimize extravasation even in organs with discontinuous capillaries). Infliximab concentrations were determined within the same workflow as the concentrations of the surface charge variants (see above and Supplemental Figure 2). The pooled infliximab serum data from all time points and animals contained two (out of 15) measurements, which were identified as outliers by a robust outlier detection algorithm (ROUT with an upper false discovery rate limit of Q=0.5%, [34]) and thus excluded from analysis. The fractional contribution (*f*_res_) of residual plasma (with the concentration *c*_Infliximab_(Serum)) to the total tissue concentration (*c*_Infliximab_(Tissue)) was then calculated according to

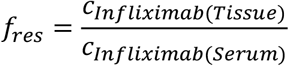

for each tissue. Based on these correction factors, the total tissue (*c*_Test substance_(Tissue)) and plasma (*c*_Test substance_(Serum)) concentration, the tissue concentration corrected for the residual plasma contribution (*c*_Test substance, corr_(Tissue)) for a given antibody variant at each time point will be

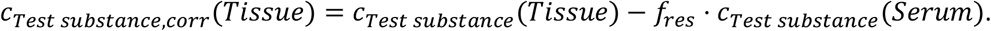

Note that due to slight differences in sampling times, serum values at t=5.5 h were used to correct the t=6 h tissue levels. Similarly, the tissue exposure corrected for the residual plasma contribution (*AUC*_corr_) was obtained according to

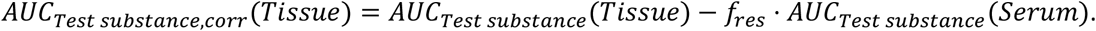

For tissue interstitial fluids, the same approach as for tissue was applied to correct for the plasma fraction in interstitial fluid.

#### Tissue-to-serum and AUC ratios

Area under the curve (AUC) ratios and the associated variances were calculated using the AUC corrected for residual serum contribution (see above) and considering the propagation of uncertainty [35]. Tissue levels from our biodistribution study in BALB/c nude mice, and serum levels obtained from the study in CL57/BL6 mice with a very similar design and sampling scheme, were used for these analyses. Note that due to slight differences in sampling times, serum values at t=5.5 h were used to correct the t=6 h tissue levels.

#### Correlation analyses and figure preparation

Correlation analyses were conducted and figures were prepared in Prism 8.4.2 (GraphPad, LLC), and illustrations were created with biorender.com.

### In vitro Transwell experiments

Experimental conditions routinely used in Transwell experiments to test for endothelial integrity were applied to establish a transendothelial transport model. Twenty-four well Transwell inserts (Costar 3470-Clear) were coated apically with 200 µl gelatine (0.1%) at 37°C for 15 min. Two hundred microliters of human microvascular endothelial cells suspension (Lonza, Basel) (6.25×10^5^ cells/ml) were seeded on the apical side of the Transwell membrane while 800 µL of medium was added on the basal side (Lonza, Clonetics EBM-2, cat. no. CC-3156; Supplements: Clonetics EGM2_MV SingleQuots CC-414). Twenty-four h post seeding, the baseline trans-endothelial electrical resistance (TEER) was measured and cells were incubated with the various antibodies (100 µM) in presence or absence of tumor necrosis factor (TNF)-α(30 ng/ml, Gibco, cat. no. PHC3016) for 24 h. TEER was measured again and antibody extravasation through the endothelial barrier was quantified by enzyme-linked immunosorbent assay (ELISA) (Abcam, cat. no. ab157709) according to the manufacturer’s instructions.

## Results

### Charge patches affect tumor uptake and biodistribution

To investigate the impact of charge patches on in vivo tumor uptake and distribution into selected tissues, we employed variants of an IgG1 antibody, which was derived from an anti-CD44 antibody and developed as a non-binding antibody, i.e. it had no detectable affinity to a specific target. Starting from this common parent molecule, three variants were engineered, carrying either negative patches (*AbNeg*), a balanced charge distribution (*AbBal*), or positive patches (*AbPos*) on their surface (Figure 1b,c). Histidines were not allowed in generation of the positive patches to avoid changes in protonation state in the physiologically relevant pH range. As an additional measure to exclude any target-related effects, the xenograft model employed (see below) does not express the antigen (CD44) in significant quantities.

To quantify interactions with negative charge-bearing surfaces such as the endothelial glycocalyx and cell membranes, we used heparin retention chromatography, an established method for this purpose [19]. We found that the selected engineered variants covered a wide range of charge properties, as reflected in the large difference in relative heparin retention (Figure 1, Supplemental Table I). In line with previous results [17–19], the variant carrying positive patches showed significantly increased clearance in non-tumor bearing C57/BL6 mice as indicated by a markedly lower AUC 0-96h (Figure 2, Supplemental Table I). This effect is likely not due to interference with FcRn binding or release, as for all three variants the retention in FcRn affinity chromatography was well within a range that indicates efficient FcRn salvage (Supplemental Table I, [19]).

To investigate the impact of the charge patches on tumor disposition and biodistribution, we chose a xenograft tumor model derived from Calu-3 in BALB/c nude mice. All three antibodies were administered in a mixture, i.e. cassette-dosing scheme, and tissue levels were subsequently analyzed in tumor and selected organs by a LC-MS workflow (Figure 1d). This enabled simultaneous quantification and internal referencing of antibody concentrations with a minimal number of animals. In organs with discontinuous capillaries (liver and spleen), the antibody with negative charge patches (*AbNeg*) and those with a balanced surface charge distribution (*AbBal*) showed very similar disposition (Figure 3). In contrast, the antibody with positive patches (*AbPos*) showed significantly elevated levels in the first few hours in these organs. This supports the notion that increased binding to liver endothelial cells [6,19] facilitates the high plasma clearance of antibodies with positive charge patches. On the other hand, the concentrations in muscle were significantly lower for AbPos compared to AbNeg and AbBal.

Surprisingly, the overall tumor levels were quite similar for all three antibody variants. This was rather unexpected based on the reduced plasma exposure of AbPos (Figure 2, Supplemental Table I). While AbNeg and AbBal reached the maximum concentration after 6 h, AbPos showed a further increase in tumor at 24 h. This observation implies that the equilibration of AbPos between serum and tissue is slower than for the other antibodies, with net tumor uptake potentially continuing beyond the tissue distribution phase.

**2.**
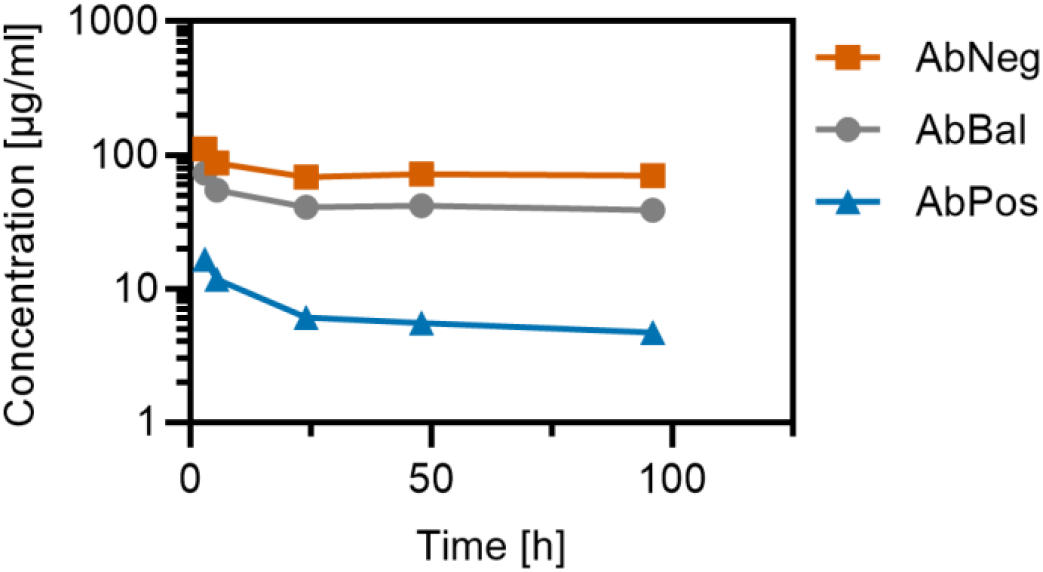
Average serum concentration-time curves of the investigated charge-patch antibody variants following single intravenous administration to tumor-bearing C57BL/6 mice as a dosing cassette at a dose of 5 mg/kg each (n=3/time point)

**3.**
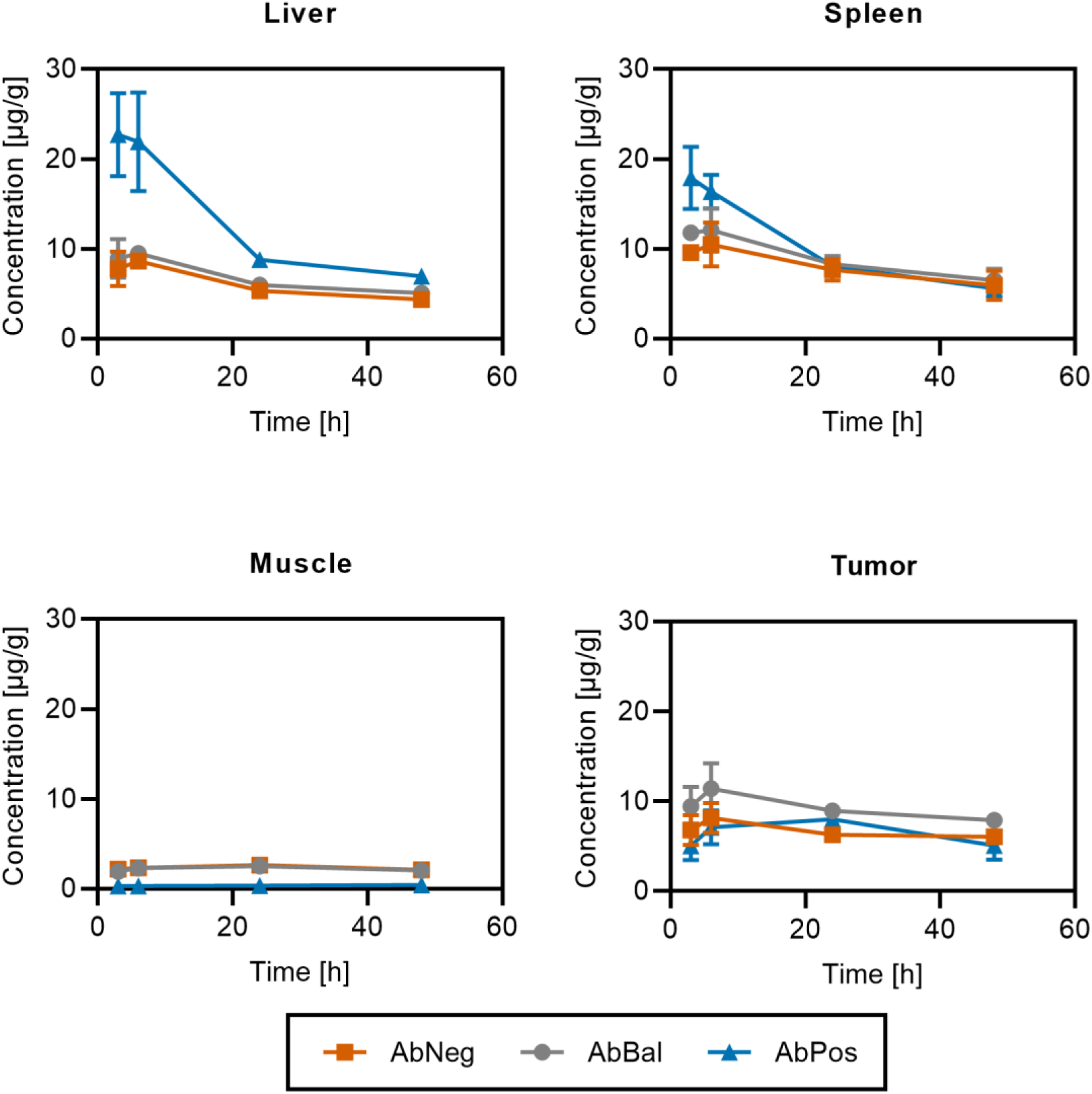
Biodistribution and tumor uptake of charge-patch variants: Shown are the tissue concentrations (mean ± SD) without residual plasma correction of all three test compounds following single intravenous administration to tumor-bearing Balb/c nude mice as a dosing cassette at a dose of 5 mg/kg each (n=3/time point).

The AUC was calculated for each of the organs, and the obtained exposures were plotted against relative retention in heparin chromatography (Figure 4). Liver exposure in the observed time range was elevated approximately 2-fold for AbPos, while muscle exposure was reduced approximately 6-fold. Tumor exposure of AbPos and AbNeg was very similar, but maximal with the balanced variant AbBal, suggesting that a balanced charged variant optimally combines tumor uptake with a long plasma half-life.

**4.**
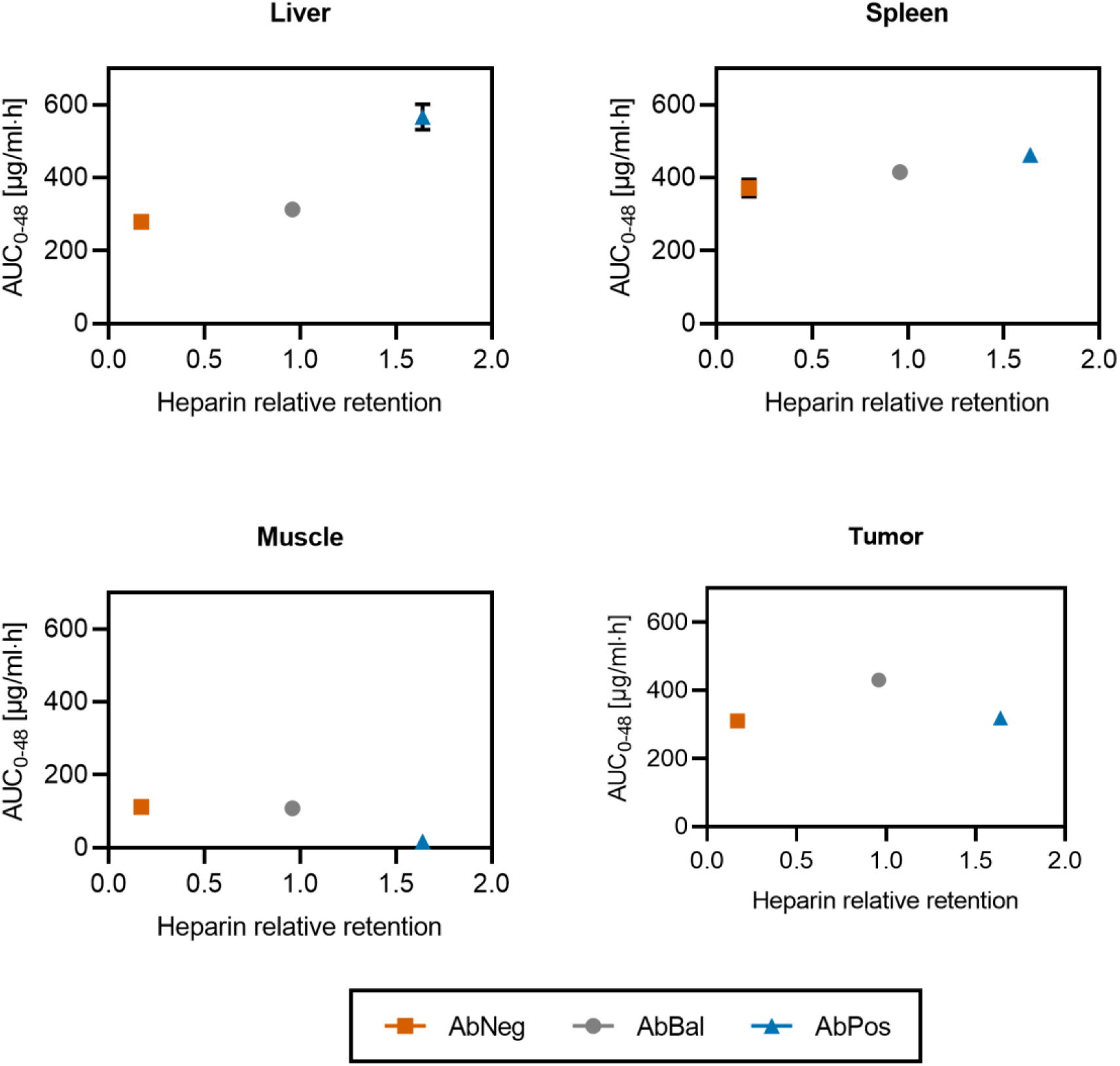
Positive charge patches increase liver exposure and reduce muscle uptake, while a balanced surface charge distribution maximizes tumor exposure. Mean AUC_0-48h_ values following intravenous dosing at 5 mg/kg each in tumor-bearing Balb/c nude mice in organs and tumor were plotted against relative retention on a heparin column (interaction with negative charge). Note that the raw exposures, without correction for residual serum, are plotted here.

### Positive-patch variant shows tissue-specific enrichment

To resolve whether individual antibody variants may show an enhanced accumulation in some tissues, we calculated ABCs [30], which are tissue-specific partition coefficients of each molecule. To exclude any effect of differing equilibration kinetics (Figure 3), potentially resulting in a ‘hook effect’ [30], we restricted our analysis to the 48-h-time point, at which steady state had been reached for all organs. Because serum samples were not available from our biodistribution study in BALB/c mice, serum levels obtained from a pharmacokinetic (PK) study in CL57/BL6 mice with a very similar design and sampling scheme (Figure 2, Supplemental Table I) were used to calculate ABCs. ABC values of AbNeg and AbBal, as well as previously published reference data [30] were quite similar, with AbBal consistently showing slightly higher values in liver, spleen, and tumor (Figure 5). AbPos, however, reached extraordinarily high ABC values of approximately 1 for liver, spleen and tumor. In contrast, the muscle ABC of AbPos was in the range of the other antibodies and the reference data.

**5.**
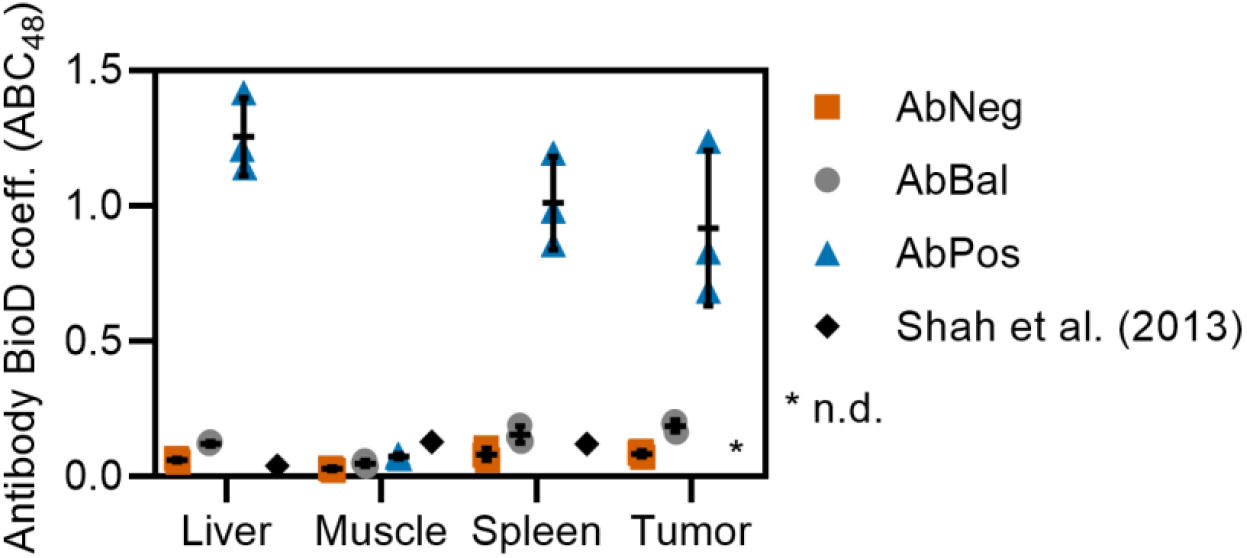
Antibody biodistribution coefficients (ABCs) at the 48-h-time point show increased partitioning of positive-patch variant AbPos into liver, spleen, and tumor tissues of tumor-bearing Balb/c nude mice. Note that large ABC values of ∼1 indicate substance-specific enrichment of the positively charged antibody (see main text). ABC values for the charge-patch antibody variants were calculated using average serum data from C57/BL6 mice, and organ and tumor data from BALB/c nude mice. Horizontal lines indicate the arithmetic mean of three animals, and error bars the standard deviation. Reference data are taken from [30].

### Positive surface patches modulate relative exposures

Having observed the enrichment of positive charge patches in some, but not all tissues, we asked how charge patches would affect the relative tissue levels (ratios of the concentrations in the respective tissue to the serum concentrations). Residual plasma can contribute significantly to the total tissue levels, particularly for well-perfused organs [30,31]. Therefore, a vascular marker (infliximab) was injected shortly before sacrifice in the biodistribution and PK studies. Amounts of infliximab, which are subsequently detected in the tissues, can then be attributed to the vascular space, with the short distribution time (5 min) ensuring that equilibration with the interstitium is minimal. We assessed the concentrations of the vascular marker in serum (Supplemental Figure 1a) and tissues (Supplemental Figure 1b), and subsequently calculated the fractional contribution of residual plasma to the total tissue concentration from their ratios (Supplemental Figure 1c). The residual plasma contribution for tumor and, in good agreement with the literature [10], for muscle, was very low (0.66%). As expected [10], residual plasma contamination was more pronounced in liver and spleen as organs with fenestrated endothelial walls (Supplemental Figure 1c). However, for these organs the tracer may reach the interstitial space within the short distribution period. Thus, the calculated fractions represent an upper limit of residual plasma contamination here. To derive tissue-to-serum ratios, the tissue levels were corrected for the residual plasma contribution. Consequently, residual plasma affected the AbNeg quantification most strongly, contributing approximately 80% of the liver exposure calculated without correction, and was negligible for AbPos (Supplemental Figure 1d). After correction of the tissue levels for the residual plasma contamination, the ratio of these to the serum levels for each molecule was calculated. The ratios were similar and quite low in muscle for all three molecules (Figure 6). However, for liver and spleen, but also tumor, tissue-to-serum ratios were markedly higher for the positively charged AbPos as compared to AbBal and AbNeg. The ratios of AbPos were essentially unaffected by residual plasma contribution, as demonstrated by repeating the analysis without the residual plasma correction (data not shown).

**6.**
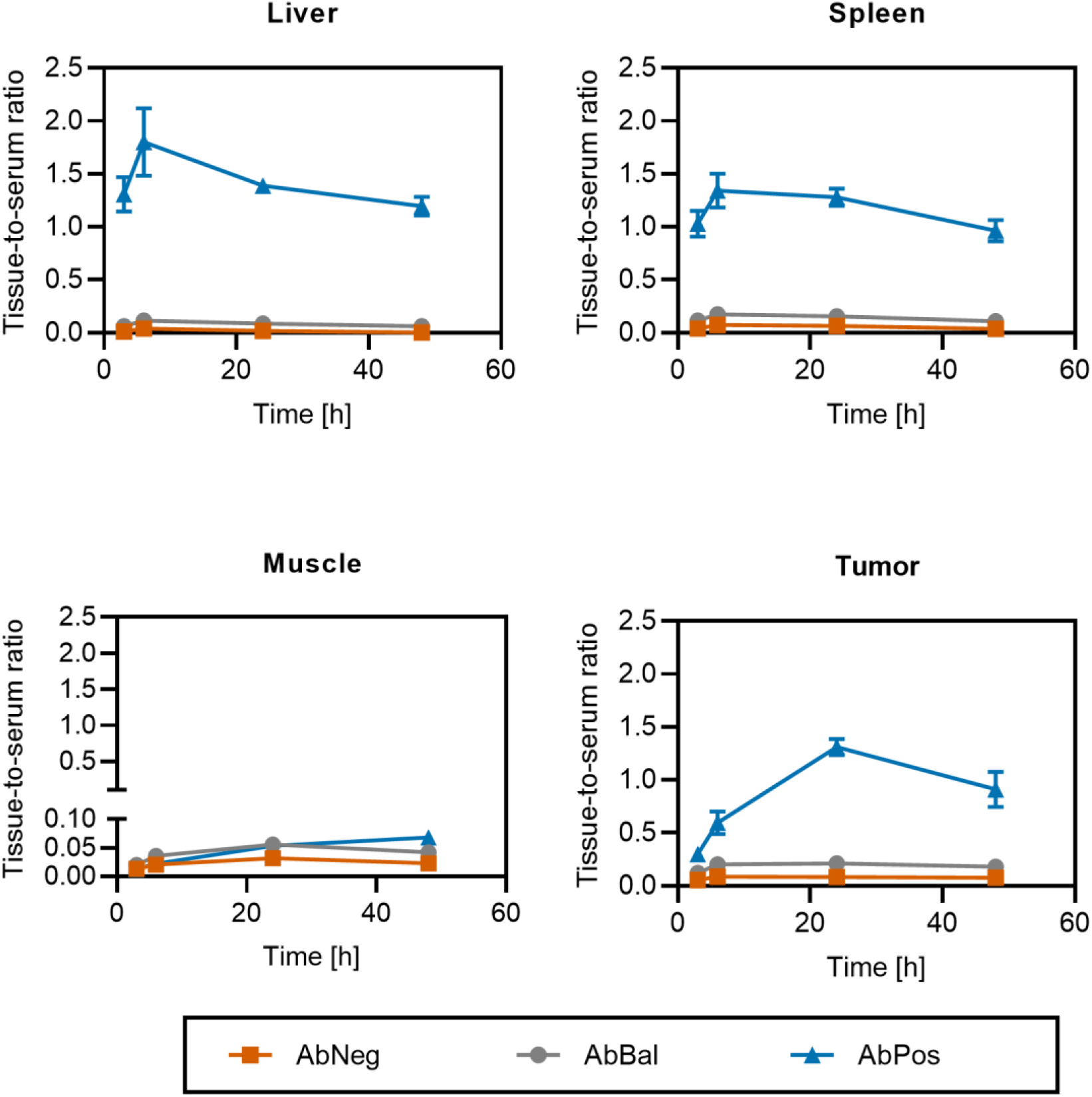
Positive charge patches in AbPos increase relative exposure in tumors and organs with discontinuous capillaries such as liver and spleen: Tissue-to-serum ratios were calculated using average serum data from C57/BL6 mice, and organ and tumor data from BALB/c nude mice (mean ± SEM, n=3/time point)

### Tissue enrichment translates to higher relative exposure of interstitial targets

We were interested in how the altered tissue and tumor levels and ratios would be reflected in the ISF concentrations. First, the ISF concentration was derived according to [36]. In agreement with the total tissue levels, interstitial levels were indeed comparable for all three antibodies in the tumor, while AbPos showed increased liver, and reduced muscle interstitial concentrations (Supplemental Figure 2). In a separate study, we used an alternative approach to experimentally determine concentrations in ISF. We employed a centrifugation protocol (Supplemental Figure 3, [31]), which enabled us to obtain (ISF and directly assess ISF concentrations applying the previously employed LC-MS workflow. The centrifugation method was used to determine ISF concentrations in muscle and skin, as ISF obtained from these tissues is less prone to intracellular fluid contamination compared to other tissues [31]. In agreement with the total tissue levels (Figure 3), the lowest interstitial concentration was observed for AbPos in both of these organs with continuous capillaries (Figure 7a,e).Using the interstitial concentrations obtained via the derivation approach [36] (Supplemental Figure 2) to calculate ISF-to-serum ratios resulted in lower values (Figure 7c). Applying the same approach for tumor (which could not be investigated using the experimental centrifugation approach) yielded significantly higher ISF-to-serum ratios (Figure 7d).

**7.**
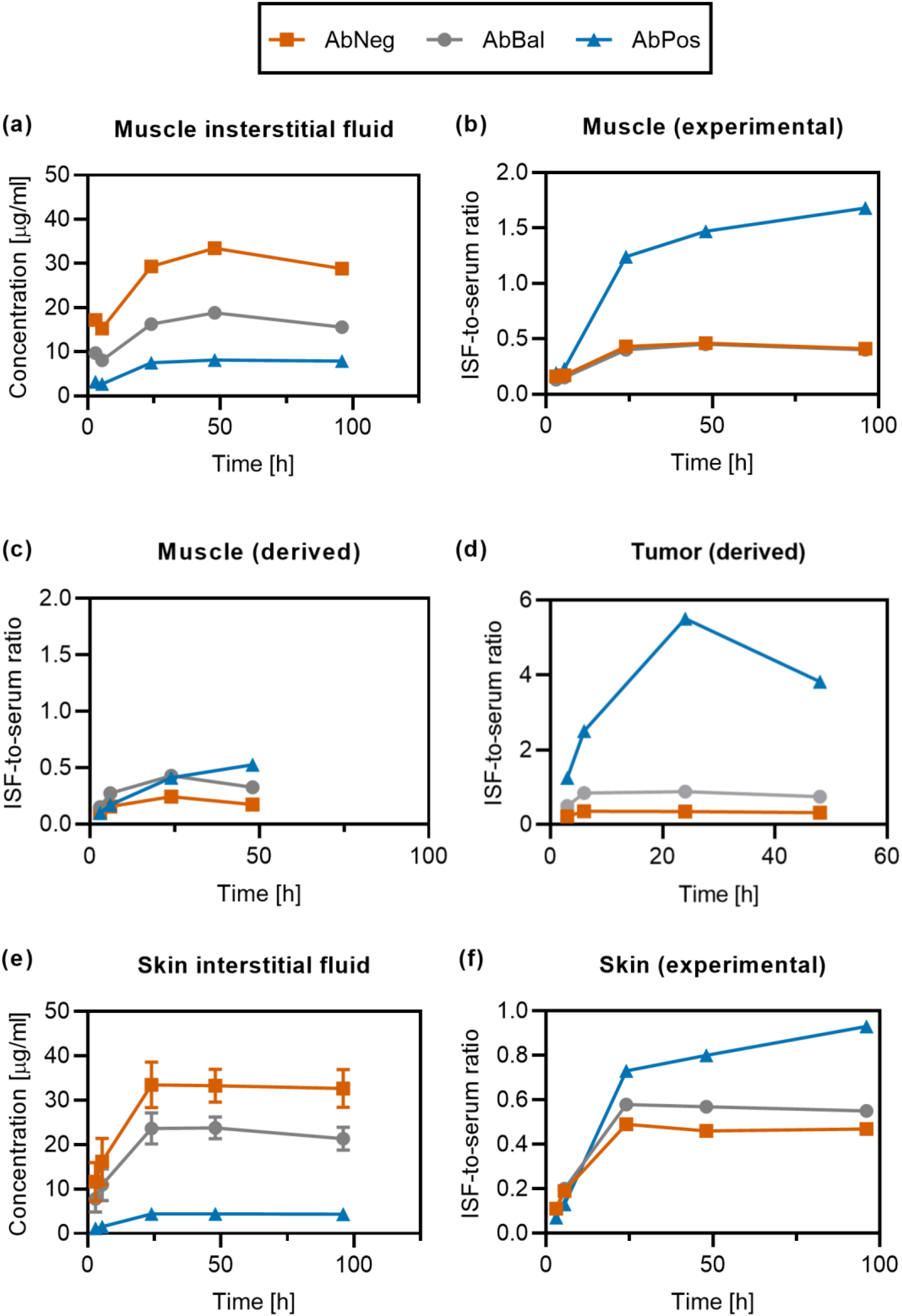
Experimental assessment of interstitial fluid (ISF) concentrations confirms relative enrichment and binding availability of positive charge-patch variant (see Supplemental Figure 3 for a schematic illustration the centrifugation to obtain interstitial fluid). (a) Antibody levels in muscle interstitial fluid in C57BL/6 mice as determined by LC-MS. (b) ISF-to-serum ratios in muscle, calculated from the experimental data shown in (a). (c) For comparison to (b), ISF-to-serum ratios in muscle were derived by the method described in [36] for the study in tumor-bearing BALB/c mice. (d) The ISF-to-serum ratio in tumor, also derived using the derivation approach of [36]. (e) Interstitial fluid levels in skin following the centrifugation protocol in C57BL/6 mice. (f) ISF-to-serum ratios calculated from the skin interstitial fluid levels as shown in (e).

While AbBal also did not reach the same absolute levels as AbNeg, we believe that this is a direct consequence of the slightly reduced serum exposure of AbBal seen in this study (Figure 2). This notion is supported by the similar ISF-to-serum ratios for AbPos and AbBal (Figure 7b,f). Their ISF-to-serum ratios increased from early time points to the 24 h time point. Thereafter, they remained similar until the last sampling time at 96 h. In contrast, the ISF-to-serum ratio for AbPos was markedly higher from 24 h post-dose onwards, and continued to increase up to the last sampling time at 96 h. In muscle, the ISF-to-serum ratio even exceeded a value of 1 from 24 h onwards.

### Transwell model reflects enhanced extravasation properties

Finally, we aimed to design an *in vitro* assay, which is predictive for the observed *in vivo* differences in tumor and organ uptake. At the same time, it should enable further insights into the underlying mechanisms, through which positive charge-patch variants are able to achieve higher tissue-to-plasma ratios as compared to ABNeg and AbBal. Therefore, an *in vitro* model scalable to high throughput, which recapitulates the limiting processes of antibody extravasation (Figure 8a) was used here. We generated a monolayer of commercially available primary endothelial cells in a conventional Transwell device, and induced tumor-like leakiness of this barrier [37] by addition of a cytokine (TNF-α). To confirm that this system is capable of passive, paracellular transport, which is the major route of tumor uptake [8] and independent of FcRn-mediated transcytosis, we used fluorescently labeled (negatively charged) dextrans (Dex-70 kDa), with a hydrodynamic radius similar to an IgG [25,38]. Indeed, induction of permeability by TNF-α lead to an increased flux to the bottom of the Transwell device (Figure 8b). The increased permeability caused by TNF-α was also reflected in reduced TEER (Figure 8c). Importantly, TEER was not reduced by the addition of Dex-70 kDa, as well as antibody variants, confirming that none of the molecules by itself induces endothelial leakiness (Figure 8c). Subsequently, the amount of antibody transported across the monolayer was quantified by ELISA for an extended set of charge-patch antibody variants, including the three molecules studied in mice. Despite the simplicity of the model system, molecules could be well differentiated based on their transendothelial transport propensity. Transendothelial transport trended with the presence of positive charge (Figure 8d), in agreement with the *in vivo* data. Of note, AbNeg showed lower transendothelial transport, while AbBal and AbPos resulted in comparable values (Fig. 7d). Importantly, the transport appeared to be independent of FcRn interactions, as indicated by experiments with Fc mutants with abolished (AbPos-3_AAA, [39,40] or elevated (AbPos-3_YTE, [39,41]) FcRn binding affinity. In summary, this method allowed to detect differences in transendothelial transport, but direct correlation with heparin binding was only moderate.

**8.**
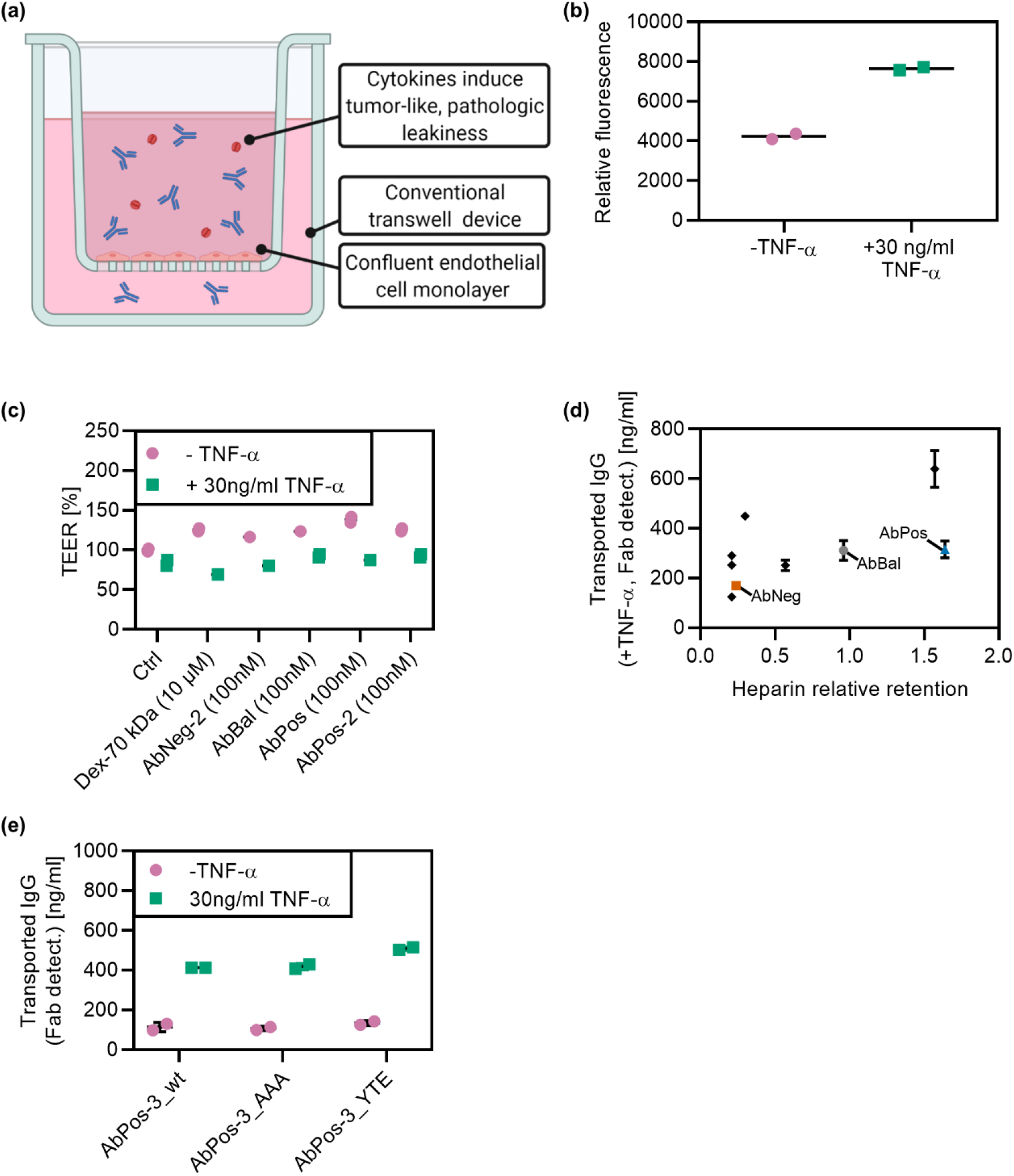
Transwell-based extravasation assay predicts in-vivo extravasation behavior. (a) Transwell experiment design modelling tumor extravasation. (b) Control experiments with fluorescently labeled dextrans with the hydrodynamic radius of an IgG show enhanced transport in the presence of TNF-α, suggesting passive paracellular diffusion as an important transport route. (c) As expected, transendothelial electrical resistance (TEER) is reduced by TNF-α, but is not significantly affected by the presence of antibodies. (d) In agreement with the behavior observed in vivo, transendothelial transport in presence of TNF-α trends (Spearman’s ρ=0.6263; p=0.0712 (two-tailed)) with the presence of positive charge patches. Black symbols represent additional charge-patch variants, engineered in the same manner as AbNeg, AbBal and AbPos. (e) Transport across the endothelial barrier seems to be independent of FcRn binding, since it is neither changed by FcRn nonbinding (AAA) nor binding-enhanced (YTE) mutations. AbPos-3 wild-type data shown here are also included in panel (e). Horizontal lines indicate the arithmetic mean of duplicates, and error bars the standard deviation.

## Discussion

This study explored the impact of distinct charge patches on the surface of therapeutic proteins on their tumor uptake and biodistribution. To this end, we deliberately engineered such patches into the Fv region of a therapeutic antibody. Importantly, this leaves the Fc region and binding to cognate receptors unaffected (see below). This is also in contrast to previous approaches, which typically employed non-selective chemical modifications leading to cationization, anionization or neutralization [17]. Chemical modifications, which are often stochastic in nature and difficult to control, would add additional complexity in manufacturing for therapeutic applications in humans. In addition, non-natural chemical modifications may give rise to safety-related concerns [17]. Using this protein engineering approach, we were able to create variants with widely different charge distribution profiles based on a single common ancestor molecule.

In the study, we used various approaches to assess biodistribution of mAb charge-patch variants. Besides the analysis of drug levels in tissues, we assessed the penetration in tissue interstitial fluid obtained by the centrifugation method. The in vivo studies were furthermore complemented by in vitro experiments, using a cell monolayer model to mimic the extravasation step of tissue uptake. While the role of extravasation in the tissue uptake of large molecules is well-known [2,8], we were interested whether in vivo differences in tissue uptake can be mirrored in the in vitro model. In this case, such an in vitro model would deserve further characterization in order to use it in the long run as replacement for tissue uptake studies in animals.

In both in vivo studies, all three test compounds were administered by cassette dosing, with subsequent specific analysis in biological samples by LC-MS. This approach reduced the number of required test animals. In addition, the cassette dosing is expected to reduce the contribution of interindividual variability in the comparison across compounds. The specific analysis included infliximab, which was used as a vascular marker. Infliximab was chosen as a commercially available antibody with a marker peptide that does not interfere with the LC-MS analytics of the test compounds. Dosing of a monoclonal antibody shortly before sacrifice as a vascular marker was used previously [31]. In the present study, we shortened the time between dosing and sacrifice from 10 min in ref. 31 to 5 min to reduce unwanted extravasation of the marker.

Increases in volume of distribution, maximum concentration in liver and other organs, and distribution kinetics have been reported as a result of introduction of positive charges through rather harsh modifications [17]. Here, we now found surprisingly marked differences in tissue uptake (Figure 3, Figure 4) solely on the basis of protein engineering. Literature suggests that the change in isoelectric point (pI) needs to be greater than 1 pI unit in order to result in a significant effect on biodistribution [17]. However, the pI difference between AbBal and AbPos was negligible (Supplemental Table I). Together with the restriction of our modifications to the Fv portion of the antibody, this highlights that localized charges, i.e charge patches, can serve as a more fine-grained metric than total antibody pI because charges are effectively screened on a length scale significantly smaller than an IgG [42]. Of note, the total tumor exposure was maximal for AbBal, for which the slightly reduced serum exposure is compensated by higher uptake.

Overall, the antibody with positive charge patches (AbPos) showed PK properties reminiscent of a smaller IgG-derived construct, such as an Fab fragment: High plasma clearance is offset by rather facile extravasation and rapid distribution into organs with discontinuous capillaries (which contribute strongly to the clearance [6]), but also uptake into the tumor tissue. By contrast, tissue levels of AbPos were lower compared to AbBal and AbNeg in both muscle and skin, as shown for both total tissue levels and interstitial fluid levels (Figures 3, Figure 7). Prompted by these findings, we wondered whether there would be an effect beyond accelerated tissue distribution for AbPos. Altered tissue equilibration kinetics can be explained by enhanced vascular permeability alone [27,43] and result in higher maximum concentration (*C*_max_) values, but may not improve steady-state concentrations or exposures, if lymph transport or cellular clearance are also elevated. Therefore, ABCs (Figure 5) and tissue-to-serum ratios (Figure 6) were calculated, which requires serum concentrations [30]. Because serum data were not available from the biodistribution study in Balb/c nude mice, we resorted to serum levels obtained from CL57/BL6 mice. Literature shows that serum clearances are comparable between most (wild-type FcRn) mouse strains, including C57BL/6, athymic nude, Balb/c, and SCID (but not NSG) mice [36,44].

ABCs are the proportionality factor between the serum concentration and concentration in the respective tissue. Hence, if mere equilibration of the tissues with the serum takes place, ABCs are (in absence of antigen binding) typically limited to a value significantly lower than 1, because only a fraction (13-24%, depending on the tissue [10,11]) of the tissue space is filled with interstitial fluid and physically accessible for the molecules. Therefore, the high ABC values (Figure 5) of AbPos in liver, spleen, and tumor indicate the enrichment in these tissues, which may result from increased transport into the organ, (specific or non-specific) binding to negatively charged tissue components such a cell surfaces or glycosaminoglycans in the extracellular matrix, cellular uptake within the tissue, or an increased accessible volume. Accordingly, the tissue-to-serum ratios of AbPos in these tissues were elevated (Figure 6). In ISF of skin and muscle, ISF-to-serum ratios increased over time to values close to or exceeding 1 (Figure 7). This increase may be due to a protracted elimination from tissue as compared to serum, potentially due to some tissue retention by charge-based unspecific binding. Wiig et al. [45] investigated the effect of an elevated pI on the steady-state levels in muscle, and found that the accessible volume was also increased in this tissue by a higher pI. However, this study compared an IgG1 with an IgG4 molecule. Using molecules derived from the same IgG (and thus, the same subclass) and maintaining a similar overall pI, we now confirm this observation. Albeit the total uptake into muscle is lower for AbPos, this is a direct consequence of the lower serum exposure and the ISF-to-serum ratios are again highest for this variant. Thus, these findings would be in line with the interpretation that interactions with negatively charged ECM components lead to some interstitial trapping of antibodies carrying positive charge patches.

Many therapeutic antibodies, in particular those directed against solid tumors, will need to reach the tissue interstitium to elicit their therapeutic effect [7,46]. Interstitial concentrations were first derived assuming a constant interstitial volume fraction. However, application of this derivation approach [36] requires the assumption that the accessible volume fraction is independent of antibody properties and the tracer molecule used, and that there is no (increased) endothelial cell binding or adsorptive endocytosis. In our case, with antibodies with strongly modified surface charge distribution, accessible volume and cell binding/uptake may vary [19,29]. Therefore, we employed a centrifugation protocol to directly assess interstitial concentrations as relevant concentration for pharmacological effects in tissues [29,31]. The increase that can be expected based on increasing the relative available volume has previously been determined by elaborate steady-state measurements, and is <30% for a cationized IgG [29] in skin. In skin, we observed an increase in ISF-to-serum ratios that was significantly higher (∼70%, Figure 7f). In muscle, even a >400% increase for AbPos (Figure 7b) was observed, corroborating that this variant indeed also preferentially localizes to the interstitial space of tumor (Figure 7d) and other tissues with discontinuous capillaries. The experimental ISF-to-serum ratios obtained through centrifugation are even higher than the derived values. This is consistent with previous reports, which also showed that the derivation approach, in turn, results in higher values than assessment by microdialysis [31,36,47]. Unfortunately, such data are not available for tumors, but the good correlation between calculated and measured interstitial concentrations for muscle (Figure 7a,b) inspires confidence that the elevated relative levels of AbPos in tumor are not due to nonspecific (endothelial) cell binding or adsorptive endocytosis. Thus, the derived ISF-to-serum ratios for tumor (Figure 7d), which are about 10-fold higher than for muscle, imply that positive patches indeed benefit the exposure of interstitial tumor targets as compared to the serum.

Extravasation has been identified as the rate-limiting step of antibody tumor disposition [8]. In line with these analyses, we found no differences in interstitial diffusion, modelled by fluorescence recovery after photobleaching (FRAP) and fluorescence correlation spectroscopy (FCS) experiments in extracellular matrix, in early exploratory experiments (data not shown). Therefore, we explored an *in vitro* assay, which should be predictive of the *in vivo* properties and, at the same time, enable mechanistic insights into the tumor extravasation process. Reducing the system to the essential components, we employed a simple Transwell model, in which the permeability of an endothelial monolayer was increased through addition of TNF-α to obtain a tumor-like, leaky endothelial barrier. The interstitial pressure is assumed to be elevated in tumors [48], lowering the effective transcapillary transport. This aspect is not captured by our model; however, it is unlikely that this bulk fluid effect differentially impacts antibody charge variants. Nonetheless, it is remarkable that the model was able to differentiate the transport propensity of a series of antibody variants with good sensitivity. Previous work revealed an association of elevated FcRn-mediated transcytosis with higher clearance, which was in turn increased by positive charge [49]. Importantly, modulation of FcRn binding alone does not result in altered uptake into liver or spleen, for example [4,5]. The experiments with FcRn binding mutants now suggest that transcapillary transport may be independent of FcRn interaction. While the correlation of transport in the Transwell assay with heparin retention was only moderate, this does not exclude a higher predictivity of the *in vivo* situation. Thus, *in vivo* data for more charge variants would be needed to conclusively assess whether the complexity of the tumor uptake processes *in vivo* can be sufficiently recapitulated by the proposed reductionistic *in vitro* model.

We used a xenograft model derived from the lung cancer cell line Calu-3. This model was chosen because the tumors are typically well vascularized, and do not show substantial necrosis in absence of treatment [50]. Nonetheless, it would be worthwhile to evaluate another model, and explore which impact different degrees of vascularization may have. The complexity of therapeutic antibody formats continues to increase, and these molecules may substantially differ in their pharmacokinetic and biodistribution properties based on molecular size and geometry, altered nonspecific or target-mediated clearance, or FcRn interactions [51]. Thus, it would be interesting to assess whether our findings fully apply to non-canonical antibody formats as well. Finally, a study with molecules with antigen binding and biological activity in the tumor could provide direct proof that the higher interstitial concentration we have observed also translates to an increased pharmacodynamic effect.

In summary, AbPos has a markedly shifted exposure profile, because it shows initially high early tissue levels in liver, spleen and tumors, despite reduced plasma exposure. Tissue-to-serum and ISF-to-serum ratios are high and exceed even unity in some cases (Figure 7). Strong tumor enrichment as observed by us has been described for cationized liposomes [52], but, to our knowledge, not for antibodies, which have only been modified on the sequence level. Antibodies with physicochemical properties as AbPos may be well suited for tissue targets, where high, but transient exposure is required. Remarkably, AbBal showed the highest absolute tumor exposure. Thus, the molecule with the lowest plasma clearance may, depending on the target localization, not always be the most suitable candidate. Therefore, *in vitro* tools like the one proposed by us may help to select fit-for-purpose molecules without the need for excessive *in vivo* work in early phases of drug discovery campaigns. Still, future studies will need to investigate the extended time course of tissue/ISF levels and whether the increased ABC ratios and tissue-to-serum ratios are sustained even beyond 48 h or 96 h, an aspect which exceeded the scope of this work.

Our findings can inform selection of the right molecule based on site of action, as well as localization of potential off-targets or on-target, off-site interactions. For example, one may increase the relative exposure of a target in liver or tumor, while reducing exposure of the same target also expressed on hematopoietic cells. We envision at least two ways how this could be practically implemented in antibody discovery and development: Either through selection of the appropriate molecule from a range of candidates based on the *in vitro* assays (heparin chromatography and the Transwell extravasation assay) or by including panning steps with polycationic substances already during in vitro selection campaigns. It should be noted, however, that positive charge patches may also increase immunogenicity, and this strategy may thus be less suited for long-term treatment. This risk is less relevant in other cases, e.g. for a single-dose or short-term treatment using potent agonists or immune modulators, when high tissue-to-plasma ratios are desired. Here, rather than being flagged for poor systemic PK, antibodies carrying positive charge patches may play out the advantages of their specific exposure profile.

## List of abbreviations

AAALAC: Association for Assessment and Accreditation of Laboratory Animal Care
ABC: Antibody biodistribution coefficients
AUC: Area under the curve
Cmax: Maximum concentration
CDR: Complementarity-determining region
ECM: Extracellular matrix
ELISA: Enzyme-linked immunosorbent assay
ESI-MS: Electrospray ionization-mass spectroscopy
FcRn: Neonatal Fc receptor
FCS: Fluorescence correlation spectroscopy
FRAP: Fluorescence recovery after photobleaching
FELASA: Federation for Laboratory Animal Science Associations
ISF: Interstitial fluid
IV: intravenous
LC-MS/MS: Liquid chromatography coupled with mass spectroscopy
MRM: Multiple reaction monitoring
PBS: Phosphate-buffered saline
pI: Isoelectric point
PK: Pharmacokinetics
TEER: Trans-endothelial electrical resistance
TFA: Trifluoroacetic acid
TNF: Tumor necrosis factor

## Declarations

### Availability of data and materials

Most data and all methods are provided the manuscript. Data not available in the manuscript will be provided by the corresponding author upon reasonable request.

### Competing interests

All authors except DKS are or were employees of F. Hoffmann-La Roche Ltd. They declare no conflict of interest.

### Funding

This work was funded by F. Hoffmann-La Roche Ltd. JCS was supported by a Roche Postdoctoral Fellowship (RPF-ID 512).

### Author contributions

All authors contributed to writing of the publication and approved the final version. JCS, KFR, MJE, HK and WFR designed the studies, TEK and HK designed the charge-patch variants, KFR, TP, TZ, and RV conducted in vitro and in vivo experiments, SMM provided bioanalytics, JCS, KFR, RV, MJE, DKS, HK, and WFR analyzed and interpreted the data.

## Acknowledgements

We acknowledge Claudia Senn and Frank Herting for support in planning and conducting the in vivo studies as well as Luca Ferrari and Denis Herzog for their support in bioanalysis.

## Supplemental Material

**Supplemental Table 1.**
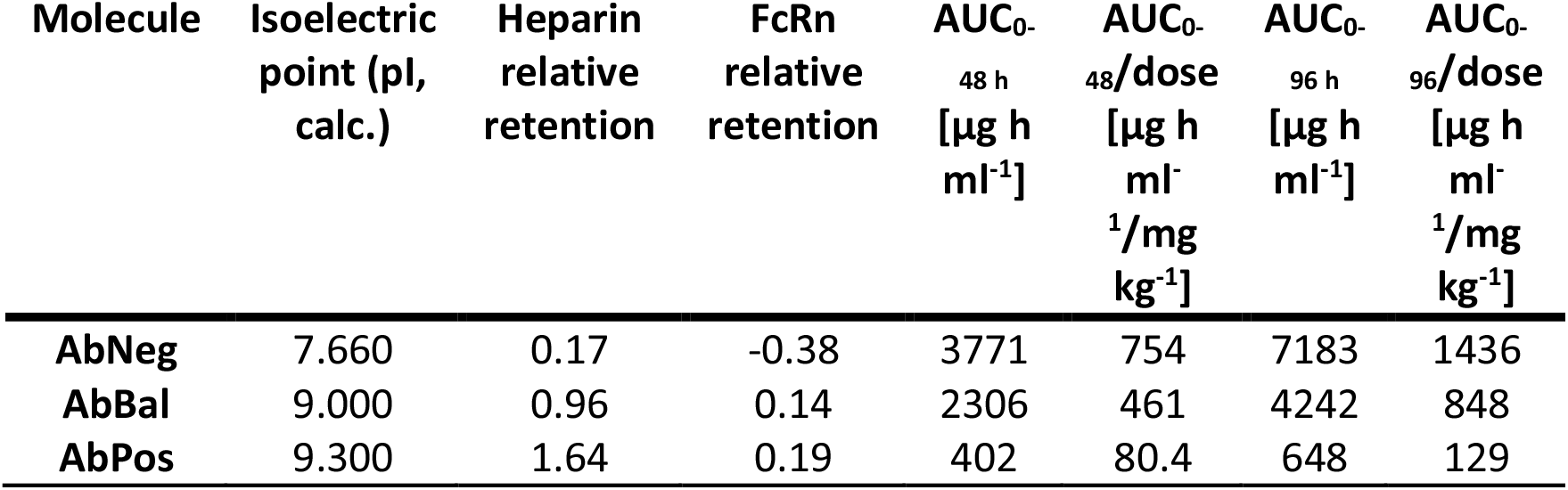
Biophysical and serum pharmacokinetic parameters in non-tumor bearing mice for charge-patch antibody variants as obtained by noncompartmental analysis.

| Molecule | Isoelectric point (pI, calc.) | Heparin relative retention | FcRn relative retention | AUC<sub>0-48 h</sub> [$\mu\text{g h ml}^{-1}$] | AUC<sub>0-48</sub>/dose [$\mu\text{g h ml}^{-1}/\text{mg kg}^{-1}$] | AUC<sub>0-96 h</sub> [$\mu\text{g h ml}^{-1}$] | AUC<sub>0-96</sub>/dose [$\mu\text{g h ml}^{-1}/\text{mg kg}^{-1}$] |
| --- | --- | --- | --- | --- | --- | --- | --- |
| **AbNeg** | 7.660 | 0.17 | -0.38 | 3771 | 754 | 7183 | 1436 |
| **AbBal** | 9.000 | 0.96 | 0.14 | 2306 | 461 | 4242 | 848 |
| **AbPos** | 9.300 | 1.64 | 0.19 | 402 | 80.4 | 648 | 129 |

**Supplemental Figure 1:**
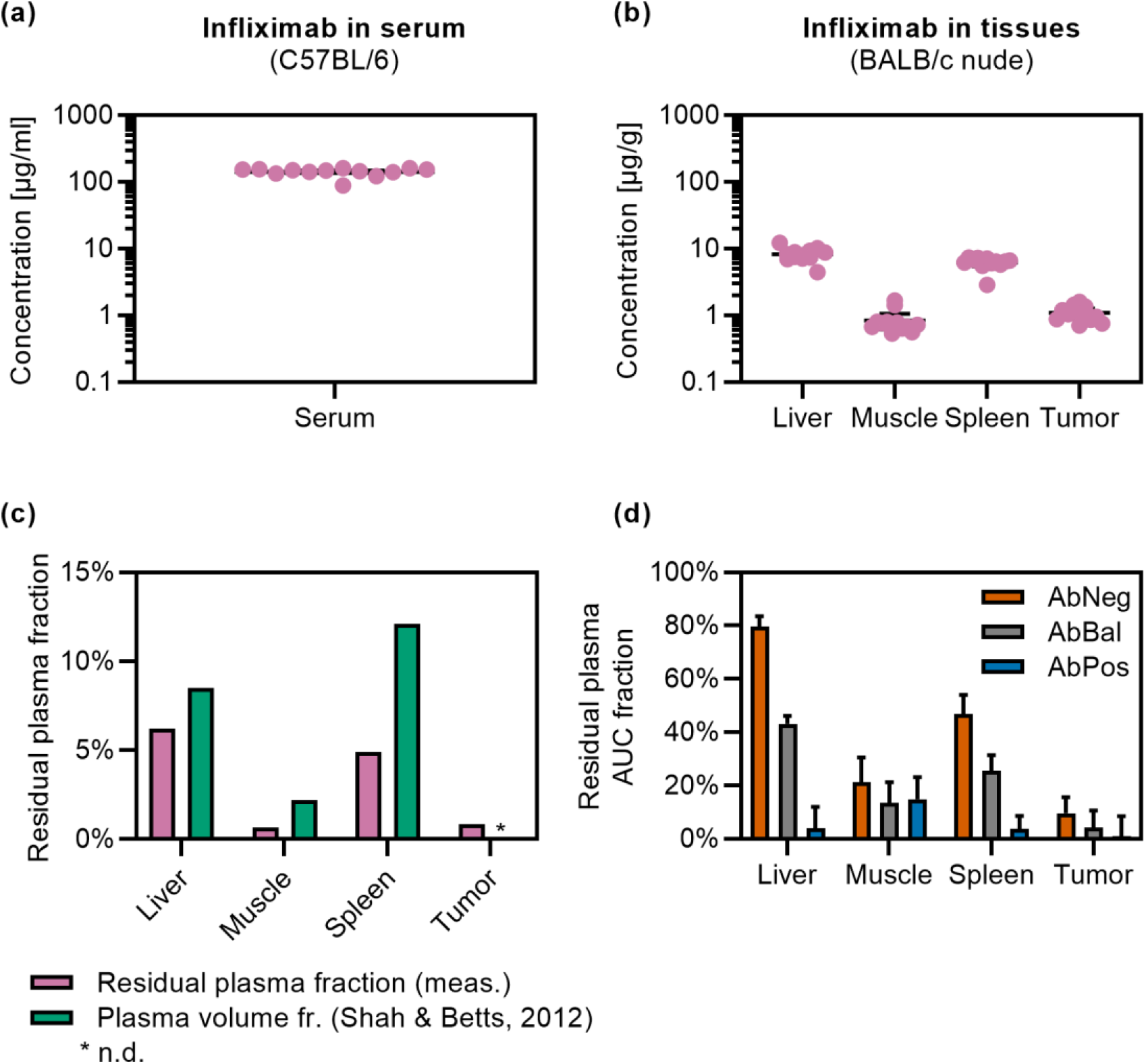
Residual plasma correction. (a-c) Levels of a vascular marker (infliximab) in serum (a) and tissues (b) were used to infer residual plasma contamination and calculate correction factors (c). Also shown is the plasma volume fraction derived from the data in [10]. (d) Relative contribution of the residual plasma fraction to the total tissue AUC in tumor-bearing Balb/c nude mice.

**Supplemental Figure 2.**
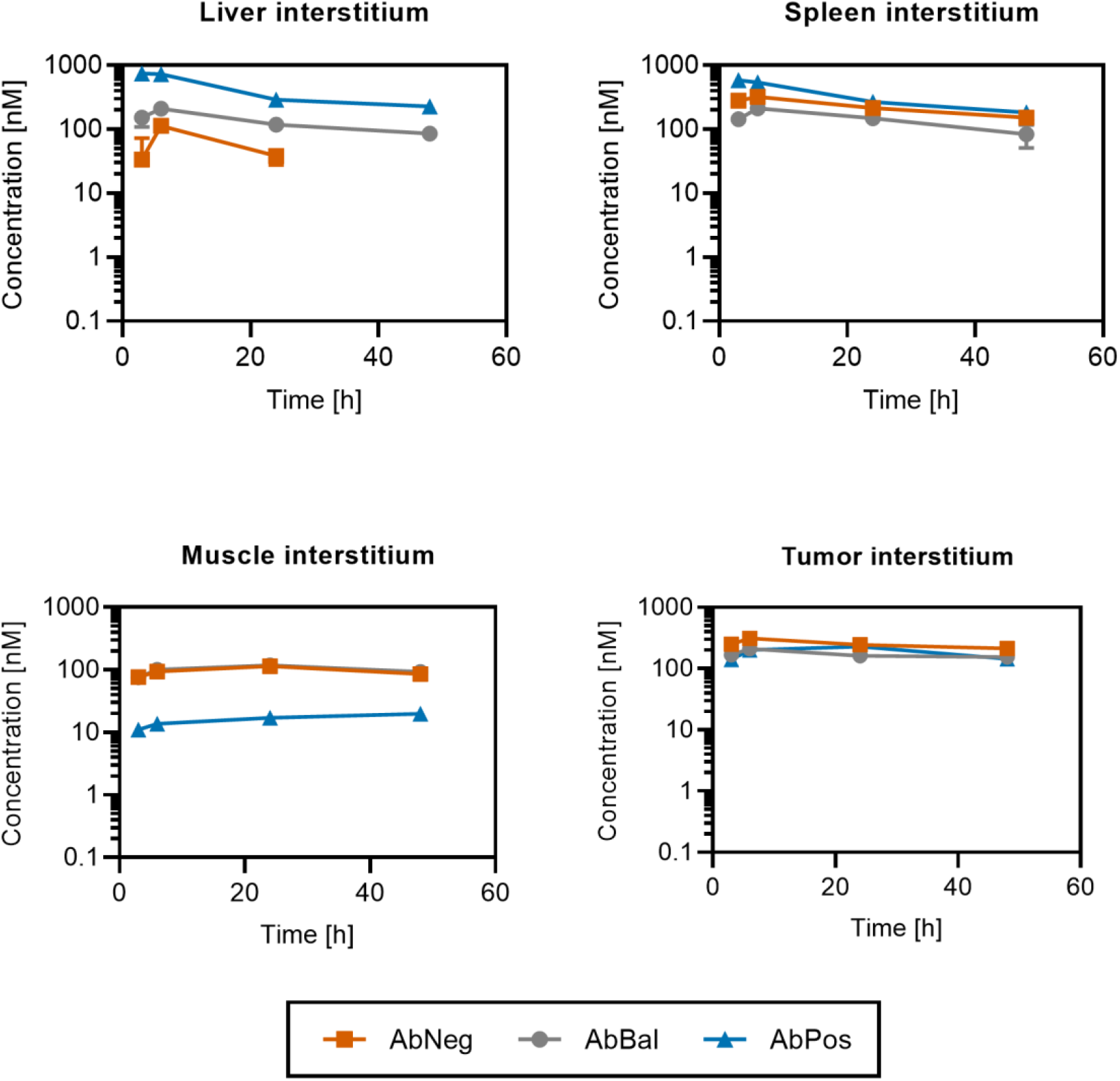
Interstitial concentrations derived from total tissue levels in tumor-bearing Balb/c nude mice using the method of Chang et al. [36] suggest similar delivery to the tumor interstitium, while liver and muscle interstitial levels differ significantly.

**Supplemental Figure 3.**
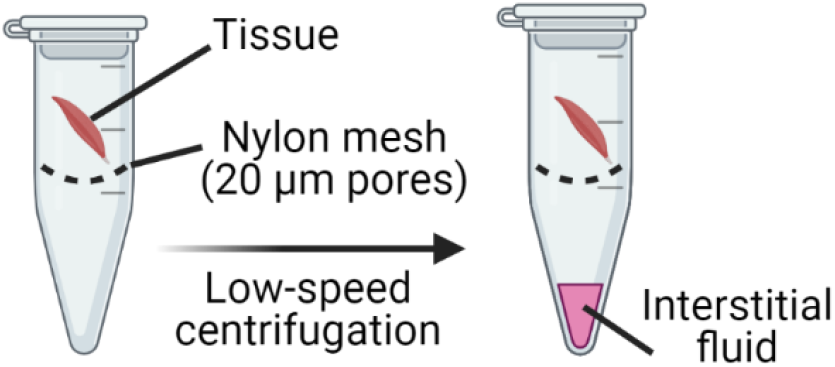
Schematic illustration of the principle of the centrifugation method to obtain interstitial fluid.

## Notes

### Summary of Updates

missing second author added

